# *RNAtor*: an Android-based application for biologists to plan RNA sequencing experiments

**DOI:** 10.1101/095315

**Authors:** Shruti Kane, Himanshu Garg, Neeraja M. Krishnan, Aditya Singh, Binay Panda

## Abstract

RNA sequencing (RNA-seq) is a powerful technology for identification of novel transcripts (coding, non-coding and splice variants), understanding of transcript structures and estimation of gene and/or allelic expression. There are specific challenges that biologists face in determining the number of replicates to use, total number of sequencing reads to generate for detecting marginally differentially expressed transcripts and the number of lanes in a sequencing flow cell to use for the production of right amount of information. Although past studies attempted answering some of these questions, there is a lack of accessible and biologist-friendly mobile applications to answer these questions. Keeping this in mind, we have developed *RNAtor*, a mobile application for Android platforms, to aid biologists in correctly designing their RNA-seq experiments. The recommendations from *RNAtor* are based on simulations and real data.

**Availability and Implementation:** The Android version of *RNAtor* is available on Google Play Store and the code from GitHub (https://github.com/binaypanda/RNAtor).

## Introduction

RNA-seq is a powerful high-throughput technology used for the understanding of complexities of transcriptomes, identification of novel coding/non-coding transcripts, spliced variants and fused transcripts, and estimation of gene and/or allelic expression. It has many advantages over the traditional low-throughput technologies like quantitative PCR or annotation-dependent methods like microarrays. However, designing RNA-seq experiments correctly, especially when prior knowledge on genome and/or transcriptome is not available, can be challenging for biologists. Knowing the numbers of replicates and sequencing reads needed to detect relative expression of transcripts accurately *a priori* are important parameters to get right biological answers. Additionally, determining the number of lanes to use in a sequencing flow cell (for example in Illumina HiSeq) and the tools to use for analyses alongside the correct experimental design will save time and money. Although some of these issues have been addressed (Busby, et al., 2013; Luo, et al., 2014), currently there is no easy-to-use, biologists-friendly mobile phone-based application (App) that can help answer these questions.

In the current study, we demonstrate the user-friendliness of an Android smartphone-based App called *RNAtor* that provides recommendations to aid biologists in the planning of the RNA-seq experiments better. The recommendations provided by *RNAtor* are based on an exhaustive combination of simulation studies and validation with real RNA-seq datasets.

## Evaluation of the App

We designed the simulation experiments to explore the number of replicates and sequencing reads requirement to draw a threshold to detect differentially expressed genes (DEGs) reliably (both in numbers and transcript recovery) at different fold changes between a control and treatment sample. Size of the transcriptome (or genome if the transcriptome size is not known) from a user-defined or from a backend database, number of replicates to use and the fold change of DEGs are user-defined parameters in *RNAtor* (Figure 1). The schema of *RNAtor* is provided in Supplementary Figure 1. *RNAtor* uses both simulated transcriptomes of various sizes (3100Mb) and a real transcriptome dataset (*Sacharomyces cerevisiae*, Accession Number ERP004763 from European Nucleotide Archive; comprising of 48 biological replicates, for two conditions; wild-type (WT) and an *snf2* knock-out (KO) mutant).

**Figure 1.**
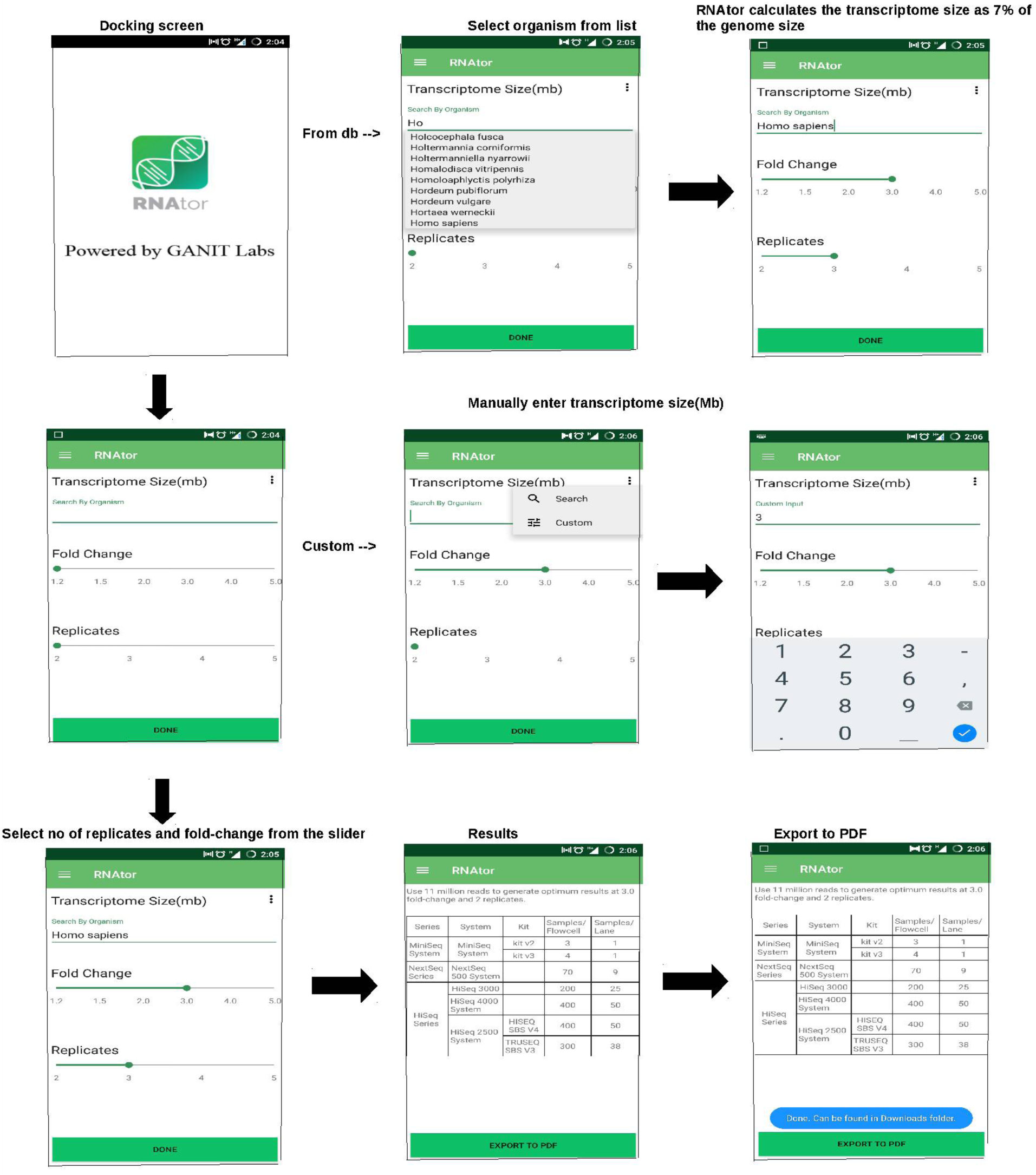
Screen shots of *RNAtor* mobile application.

*RNAtor* was evaluated using questions that a biologist typically may ask before starting an experiment followed by the recommendations provided by *RNAtor*.

**Question 1:** How many sequencing reads are needed to detect optimal number of differentially expressed genes (DEGs) at 1.2 - 5fold change for a 3Mb transcriptome with 3 replicate samples?

***RNAtor:*** For a 3MB transcriptome with 3 replicates, the following numbers of sequencing reads are needed for detecting transcript changes between 1.2 - 5 folds.

2 Millions reads for 5-fold

6 Millions reads for 4-fold

10 Millions reads for 3-fold

14 Millions reads for 2-fold

20 Millions reads for 1.5-fold

30 Millions reads for 1.2-fold

**Backend** We simulated a range of Illumina-like RNA-seq reads (0.2 - 20 millions), for human chromosome 14 (~3Mb) using Polyester (Frazee, et al., 2015). We observed that the numbers of detected DEGs simulated at a given fold change peaked for a certain coverage before plateauing (Figure 2) that remained valid for the real data (Figure 3) and simulated transcriptomes of larger sizes (10Mb, 30Mb and 100Mb) (Supplementary Figure 2). Increasing the number of sequencing reads increased the sensitivity of detection. The final recommendations from *RNAtor* correspond to the number of DEGs at its peak, and therefore a good compromise between sensitivity and cost of sequencing. Changing the number of replicates does change the recommendation. For example, with more number of replicates, *RNAtor* suggests to produce less number of reads to obtain the same information (Table 1).

**Figure 2:**
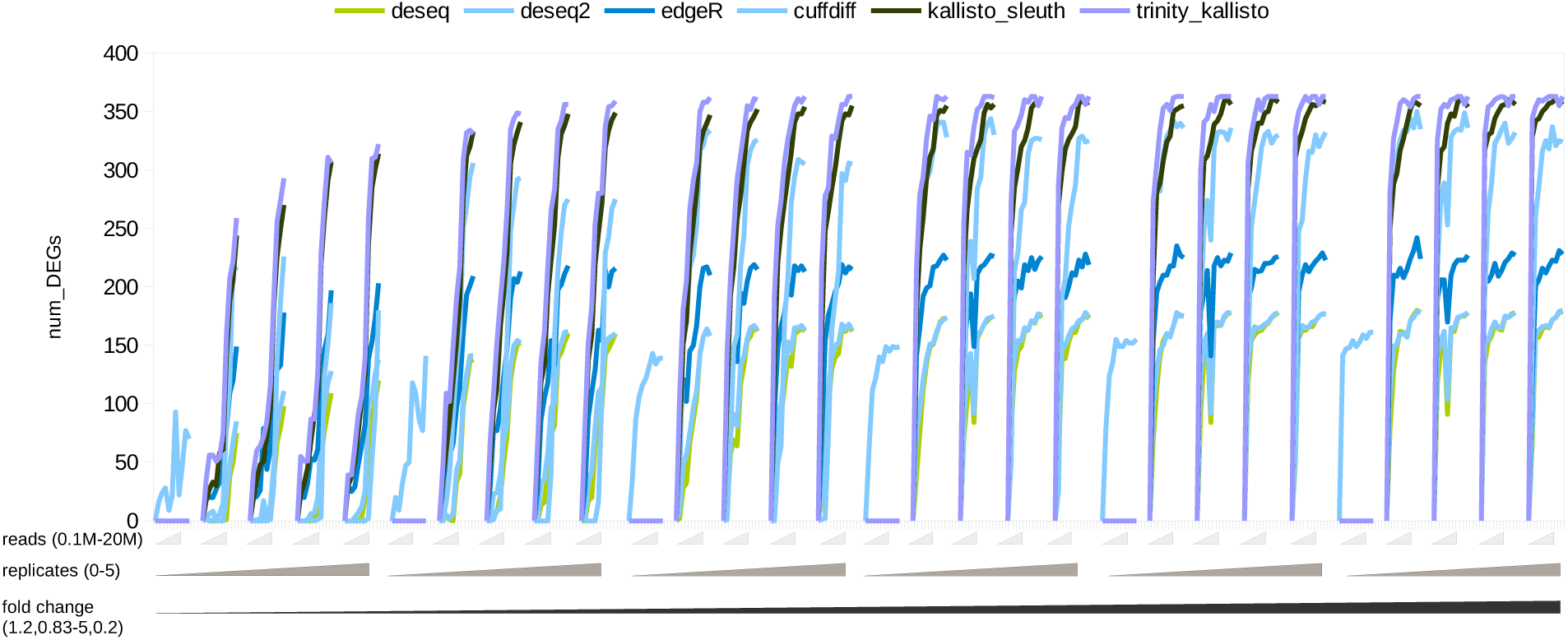
Number of differentially expressed genes (DEGs) detected for simulated data (hg19 chr14) by different tools.

**Figure 3:**
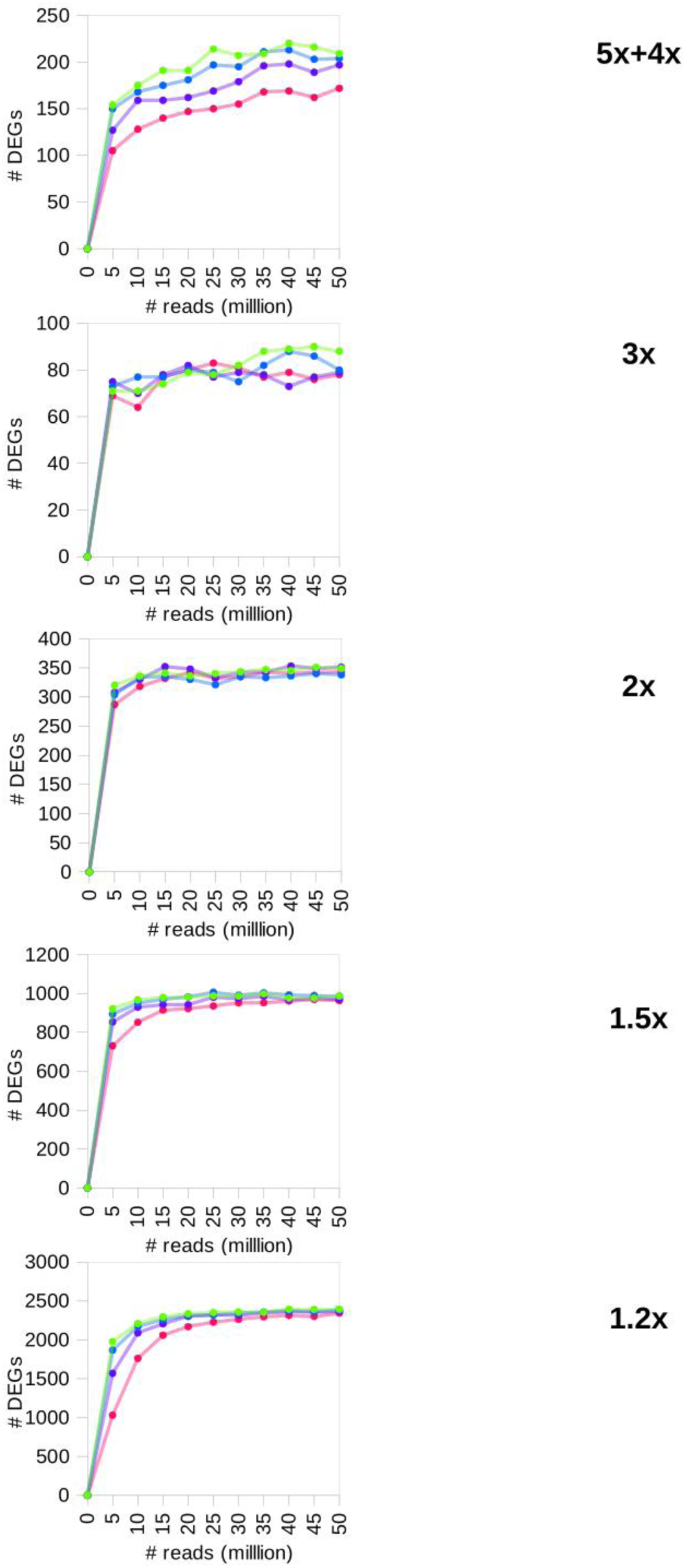
Number of differentially expressed genes (DEGs) detected for real data on *Saccharomyces cerevisiae* using the Kallisto-Sleuth pipeline.

**Table 1.** *RNAtor* output on number of sequencing reads (in millions) to be produced for a given number of sample replicates to detect differentially expressed genes at a given fold change.?

|  | 2 replicates | 3 replicates | 4 replicates | 5 replicates |
| --- | --- | --- | --- | --- |
| 5fold | 6 | 2 | 1.5 | 1.5 |
| 4fold | 10 | 6 | 2 | 1.5 |
| 3fold | 10 | 6 | 6 | 6 |
| 2fold | 14 | 10 | 10 | 6 |
| 1.5fold | 30 | 20 | 20 | 14 |

**Question 2** Which analysis tool to use in order to detect optimal number of DEGs with high sensitivity?

***RNAtor:*** Kallisto.

Backend: We compared the performance across five widely used genome-guided tools (Deseq (Anders and Huber, 2010); Deseq2 (Love, et al., 2014); EdgeR (Robinson, et al., 2010); Cuffdiff (Trapnell, et al., 2012); and Kallisto (Bray, et al., 2016) for RNA-seq data analyses. Focusing purely on the number of DEGs detected between WT and KO, Kallisto performed best over the other tools tested (Figure 2 and Supplementary Figure 3).

**Question 3:** Which genome-guided pipeline/tool to use for detection of optimal number of DEGs with high specificity and with high recovery of transcripts?

***RNAtor*** Cuffdiff for high specificity and DeSeq2 and EdgeR for high transcript recovery.

**Backend:** Although Kallisto-Sleuth is fast and produced results with high sensitivity; we observed that this was at the expense of specificity of detection (Supplementary Figure 3). Cuffdiff produced results with high specificity (Supplementary Figure 3) albeit with a loss of sensitivity (Figure 2).

**Question 4:** Out of the tools tested, which one gives a better handle on measuring transcript recovery?

***RNAtor:*** CuffDiff.

**Backend:** Transcript recovery was higher for CuffDiff at lower fold changes, especially for longer transcripts, over both DeSeq2 and EdgeR (Supplementary Figure 4).

**Question 5:** Do the above recommendations differ when using an assembly-based pipeline over the genome-guided tools?

***RNAtor:*** Not for the number of DEGs detected but the assembly-based pipeline yielded DEGs with lower specificity.

Backend: Using Trinity as an assembly pipeline (Grabherr, et al., 2011) along with Kallisto did not largely change the number of DEGs detected when compared with Kallisto-Sleuth pipeline (Figure 2). However, this happened at the cost of specificity (Supplementary Figure 3).

## Conclusions

*RNAtor* is a biologist-friendly and easy-to-use platform to design RNA-seq experiments based on certain user inputs. Where sample is limiting, *RNAtor* provides guidelines to produce required number of reads to detect differentially expressed transcripts. This can especially be useful in detecting differentially expressed transcripts at low end. Despite its usefulness, *RNAtor* has certain limitations. For example, it does not take into account, 1) the dynamic nature of transcriptome (where the exact size of transcriptome is not known and cannot simply be derived from the genome size), 2) the throughput of different sequencing instruments and 3) detection of spliced variants. These will form the basis for its future release.

## Supplementary Figure

**Supplementary Figure 1:**
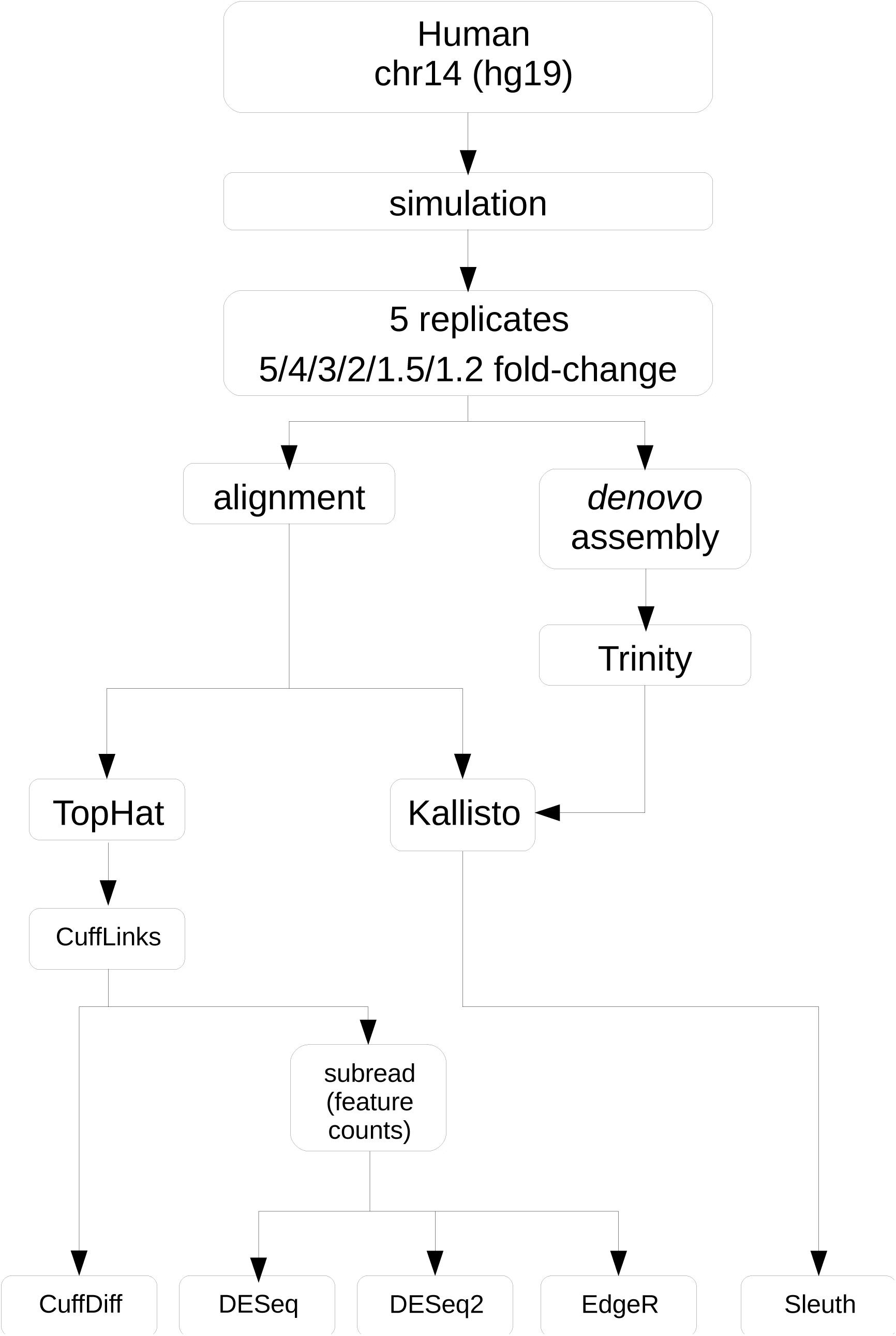
*RNAtor* schema.

**Supplementary Figure 2:**
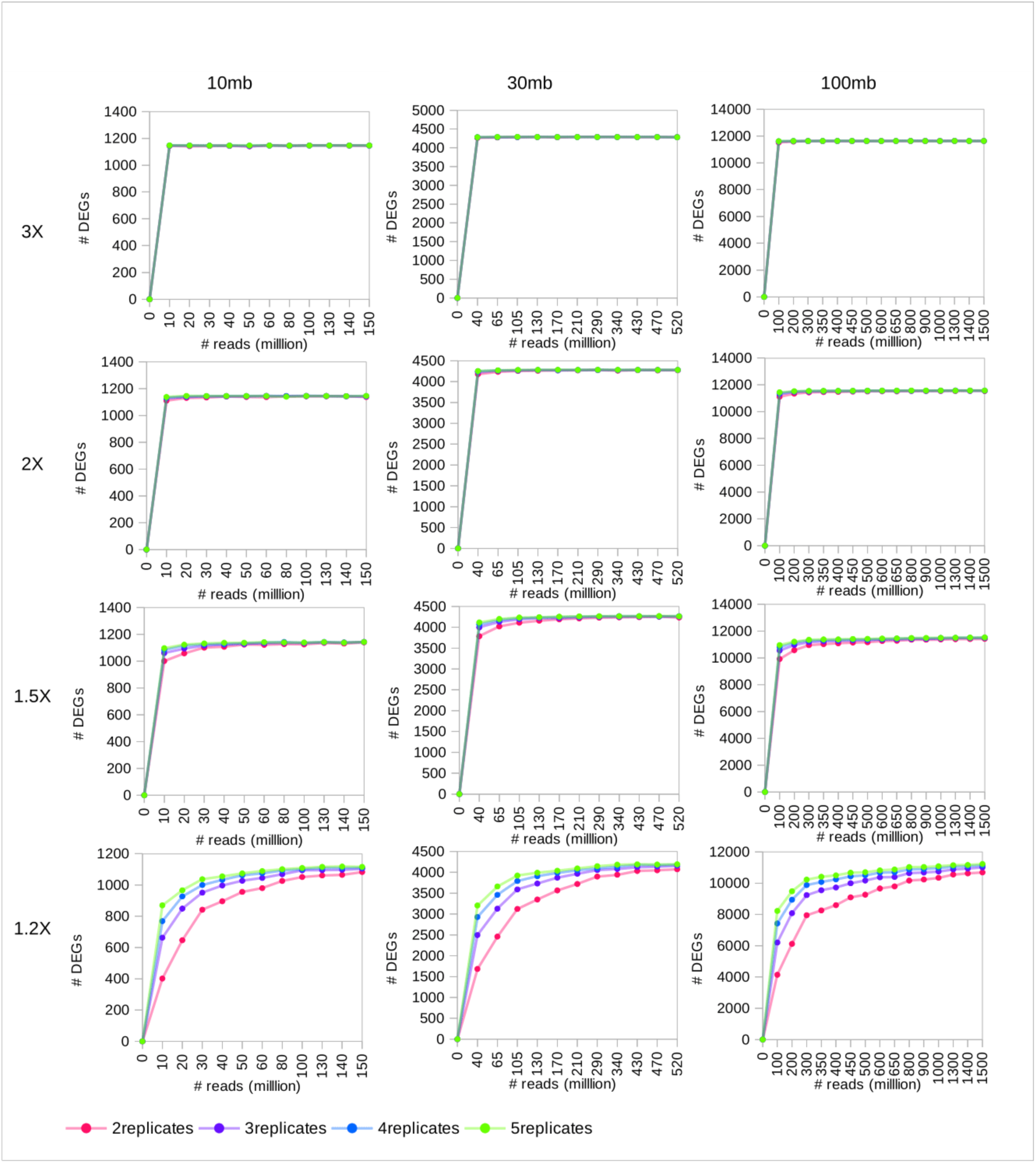
Number of differentially expressed genes (DEGs) detected for various simulated data on 10mb, 30mb and 100mb transcriptomes, using the Kallisto-Sleuth pipeline.

**Supplementary Figure 3:**
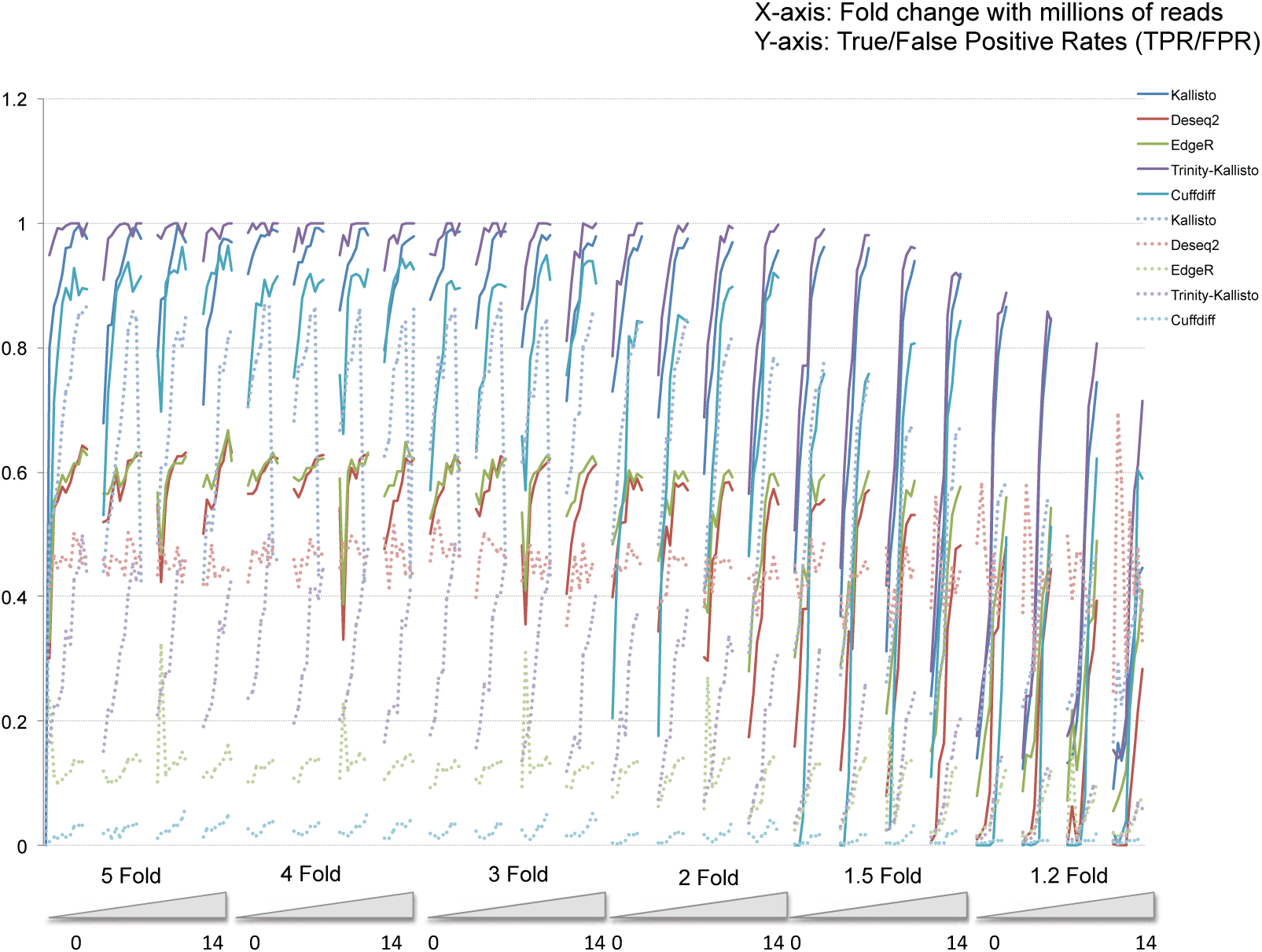
True/false positive curves for DEGs recovered under various simulation conditions by various tools.

**Supplementary Figure 4:**
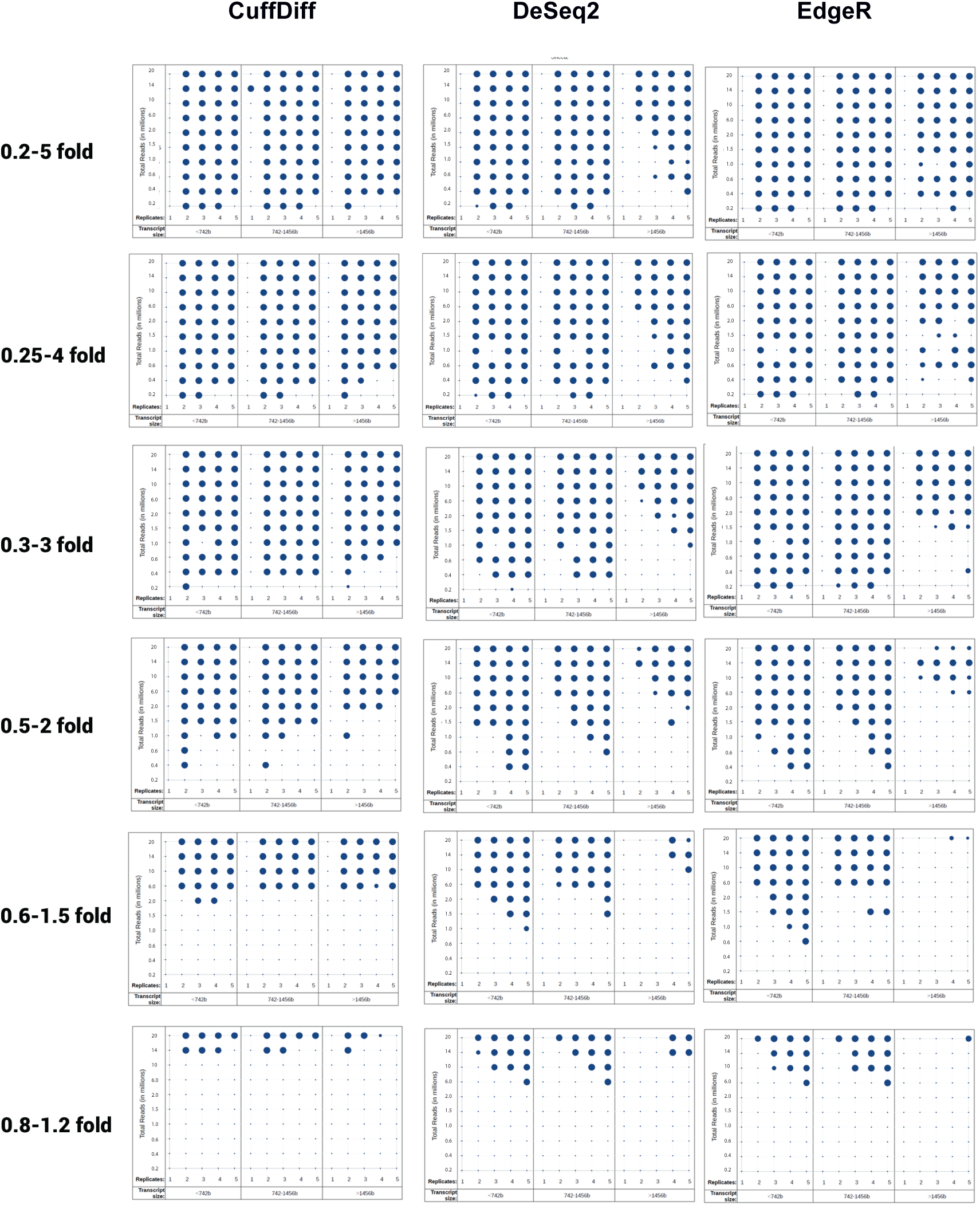
Percentage recovery of transcripts under various simulation conditions by various tools. Size of the bubble represents the extent of transcript recovery.

## References

1. Anders, S. and Huber, W. Differential expression analysis for sequence count data. Genome biology 2010;11(10):R106.

2. Bray, N.L., et al. Near-optimal probabilistic RNA-seq quantification. Nature biotechnology 2016;34(5):525-527.

3. Busby, M.A., et al. Scotty: a web tool for designing RNA-Seq experiments to measure differential gene expression. Bioinformatics 2013;29(5):656-657.

4. Frazee, A.C., et al. Polyester: simulating RNA-seq datasets with differential transcript expression. Bioinformatics 2015;31(17):2778-2784.

5. Grabherr, M.G., et al. Full-length transcriptome assembly from RNA-Seq data without a reference genome. Nature biotechnology 2011;29(7):644-652.

6. Love, M.I., Huber, W. and Anders, S. Moderated estimation of fold change and dispersion for RNA-seq data with DESeq2. Genome biology 2014;15(12):550.

7. Luo, H., et al. The importance of study design for detecting differentially abundant features in high-throughput experiments. Genome biology 2014;15(12):527.

8. Robinson, M.D., McCarthy, D.J. and Smyth, G.K. edgeR: a Bioconductor package for differential expression analysis of digital gene expression data. Bioinformatics 2010;26(1):139-140.

9. Trapnell, C., et al. Differential gene and transcript expression analysis of RNA-seq experiments with TopHat and Cufflinks. Nature protocols 2012;7(3):562-578.

